# Na^+^/H^+^ Exchanger Isoform 1 Regulates Apoptosis Susceptibility in Pulmonary Arterial Smooth Muscle from the Sugen/Hypoxia model of Pulmonary Hypertension

**DOI:** 10.1101/2025.11.06.687041

**Authors:** Manuella Ribas Andrade, Xin Yun, Michael P. Croglio, Shannon Niedermeyer, Haiyang Jiang, Nicolas Philip, Samuel Murray, Micheal Munson, Karthik Suresh, Mahendra Damarla, John Huetsch, Larissa A. Shimoda

## Abstract

Pulmonary hypertension (PH) is characterized by vascular remodeling driven in part by apoptosis-resistant pulmonary arterial smooth muscle cells (PASMCs). Na⁺/H⁺ exchanger isoform 1 (NHE1) regulates intracellular pH and plasma membrane cytoskeleton anchoring, influencing PASMC migration and proliferation, but the role of NHE1 in apoptosis remains unclear. NHE activity and NHE1 surface expression were increased in PASMCs from the Sugen/Hypoxia (SuHx) rat model compared to controls. Despite increased endoplasmic reticulum (ER) stress at baseline, SuHx PASMCs were resistant to apoptosis following H_2_O_2_ challenge. Pharmacological inhibition of NHE activity with ethyl-isopropyl amiloride (EIPA) and silencing with siRNA restored apoptosis in SuHx PASMCs. Conversely, NHE1 overexpression in control PASMCs conferred apoptosis resistance. Expression of mutant NHE1 constructs lacking ion translocation or binding to the adaptor protein, ezrin, also reduced H_2_O_2_-induced apoptosis. Mechanistically, apoptotic stimulation with H_2_O_2_ increased p38 phosphorylation in PASMCs from control, but not SuHx, rats, indicating impaired activation of this pro-apoptotic pathway. NHE1 suppression via EIPA or siRNA restored p38 phosphorylation in SuHx PASMCs, while overexpression of NHE1 (wild-type or mutants) suppressed p38 activation following apoptotic stimulation. Inhibition of p38 with SB203580 prevented the pro-apoptotic effect of EIPA, validating a role for p38 signaling in NHE1-mediated apoptosis resistance in SuHx PASMCs. These findings identify NHE1 as necessary and sufficient for PASMC apoptosis resistance in PH, by a mechanism independent of ion transport or ezrin-binding functions but involving suppression of p38 phosphorylation. Targeting NHE1-dependent pathways may restore PASMC apoptosis and offer a novel therapeutic strategy to reverse pulmonary vascular remodeling in PH.

## Introduction

Pulmonary hypertension (PH) is a complex and severe cardiopulmonary disorder defined as a mean pulmonary artery pressure >20 mmHg at rest^1^. According to the World Health Organization (WHO), PH can be clinically categorized into five subgroups according to etiology, clinical symptoms and comorbidity: (1) pulmonary arterial hypertension (PAH), (2) PH due to left-sided heart disease, (3) PH due to chronic lung disease, (4) chronic thromboembolic PH and (5) multifactorial PH^2^. In PAH, the increase in pulmonary arterial pressure is associated with sustained vasoconstriction and vascular remodeling of the distal pulmonary arteries^3,4^, the latter due in part due to increased proliferation^5–7^, migration^6–9^ and resistance to apoptosis of pulmonary arterial smooth muscle cells (PASMCs)^6,10^. Even with advancement of treatment options, there is still no cure for PH, and current therapies only alleviate symptoms^11,12^ highlighting the relevance of exploring mechanisms of vascular remodeling in PH for development of new therapeutic approaches.

The Sugen-Hypoxia (SuHx) rat model of PH combines a single exposure to SU5416, a vascular endothelial growth factor (VEGF) receptor 2/tyrosine kinase antagonist, and chronic hypoxia exposure to develop features resembling human PAH such as angio-obliterative lesions in the pulmonary arteries, severe increase in right ventricular systolic pressure, and right ventricular hypertrophy. Notably, PH induced by the SuHx model is irreversible even after return to normoxia.

Resistance to apoptosis has been reported in PASMCs from SuHx rats^10,13^ and patients^14–16^ and is believed to contribute to vascular remodeling of the pulmonary arteries. Apoptosis is the most common form of programmed cell death and plays an important role in maintaining tissue homeostasis and proper cellular turnover. In disease, cellular apoptosis serves as a protective process of quality control by eliminating cells that are abnormal or damaged^17^. However, the exact mechanisms underlying apoptosis resistance in PAH are not fully understood.

Another key feature of PH is increased endoplasmic reticulum (ER) stress in PASMC^18^, in which the ER cellular machinery responsible for protein folding becomes overloaded, leading to abnormal cell function and contributing to the vascular remodeling seen in the monocrotaline (MCT)^19,20^ and chronic hypoxia (CH) models of PH^20^. In healthy cells, increased ER stress is a common apoptotic stimulus that leads to a series of well characterized events in the ER-stress mediated apoptotic cascade. Despite evidence for ER stress, PASMCs from SuHx rats exhibit minimal apoptosis, both at baseline and in response to apoptotic stimuli^10^.

The Na^+^/H^+^ exchanger (NHE) is major contributor to maintenance of intracellular pH homeostasis^21^. In PASMCs, NHE isoform 1 (NHE1) is the main plasma membrane regulator of intracellular pH (pH_i_)^22^. In the CH model of PH, increased expression of NHE1 in PASMCs was linked to increased NHE activity and alkaline pH_i_ ^22,23^. Upregulation of NHE1 expression^24,25^ and/or activity^7^ was associated with increased PASMC migration and proliferation and development of PH. Reducing NHE activity by pharmacological inhibition^24,26^ or NHE1 silencing via knockdown or knockout was protective against hypoxia-induced PH^24,27,28^, decreasing hypoxia-induced vascular remodeling and alleviating the increased proliferation and migration in PASMCs from CH animals. In addition to its important role in intracellular pH regulation, NHE1 has also been shown to regulate migration in fibroblasts^29^ and PASMCs^25^ and PASMCs proliferation^25^ through the ezrin/radixin/moesin (ERM) protein binding motif in the cytosolic tail region of NHE1, via protein-protein interactions with phosphorylated ezrin and the actin cytoskeleton^25^. Additionally, in tubular epithelial cells, NHE1 regulated cell survival via interactions and scaffolding of the ERM proteins^30,31^; however, the role of NHE1 in regulating PASMC apoptosis has not been described. NHE1 can also function as a plasma membrane scaffold for the assembly of signaling complexes, including kinases and other regulatory molecules, through the cytosolic tail region^32^. The p38 mitogen-activated protein kinase (p38) is a well-studied signaling molecule that participates in the regulation of cellular responses to stress and plays a central role in regulating smooth muscle cell apoptosis in response to various stressors^33,34^. In lymphocytes, phosphorylation of NHE1 by p38 regulates apoptosis via increases in NHE1 activity^35^. Conversely, studies show NHE1 regulates the activity of multiple mitogen-activated protein kinases (MAPK), including p38 in fibroblasts and cardiomyocytes^36^. Whether the p38 MAPK apoptotic pathway is linked to NHE1 or plays a role in the susceptibility of PASMCs to apoptosis in the SuHx model is still unknown.

In this study, we investigated the role of NHE1 in resistance to apoptosis in PASMC isolated from the SuHx model of PH. Specifically, we performed experiments testing whether loss of function of NHE1 in PASMCs from the SuHx model increases susceptibility to apoptosis and whether increasing NHE1 in control PASMCs could induce apoptosis resistance. Our experiments also explored the mechanism by which NHE1 modulates resistance to apoptosis, focusing on the well-known role of NHE1 in pH_i_ regulation as well as non-canonical roles of NHE1 in cytoskeletal organization and p38 phosphorylation.

## Methods

All protocols were reviewed by and performed in accordance with Johns Hopkins University Animal Care and Use Committee. Protocols and procedures comply with NIH and Johns Hopkins Guidelines for the care and use of laboratory animals.

### Sugen/Hypoxia model

PH was induced in adult Wistar rats (150–200 g; Inotiv) of both sexes by a single subcutaneous injection of SU5416 (20 mg/kg), prepared in a carboxymethylcellulose (CMC) and DMSO solution as described previously^7^, followed by exposure to 10% O_2_ (hypoxia) for 3 wk (n=25). The rats were exposed to normoxic conditions for <5 min twice a week to change cages and replenish food and water. At the end of 3 wk, rats were returned to normoxia for an additional 2 wk. Normoxic control rats (Nor) were injected with vehicle and maintained in room air for 5 wk (n=25). Selection for the treatments were randomized. All animals were kept in the same room and exposed to the same 12-hr light-dark cycle. Rats were housed in standard rat cages (3 rats/cage) with free access to food and water, at ambient temperature regulated between 68-76 °F. At the end of exposures, rats were anesthetized (Ketamine, 75 mg/kg; Xylazine, 7.5 mg/kg) and depth of anesthesia was confirmed via paw pinch. Animals were euthanized via exsanguination and the heart and lungs removed and transferred to a dissecting dish filled with cold N-[2-hydroxyethyl]piperazine-N’-[2-ethanesulfonic acid] (HEPES)-buffered salt solution (HBSS) containing (in mM): 130 NaCl, 5 KCl, 1.2 MgCl_2_, 1.5 CaCl_2_, 10 HEPES, and 10 glucose, with pH adjusted to 7.2 with 5 M NaOH. Under a microscope, the atria and large conduit vessels were removed, and the right ventricle (RV) wall was carefully separated from the left ventricle and the septum (LV+S). Both portions were blotted dry and weighed.

### Isolation of Pulmonary Arterial Smooth Muscle Cells

The methods for obtaining primary cultures of rat PASMCs have been previously described^24^. Briefly, intrapulmonary arteries (PAs; 200-600 µm outer diameter) were dissected from lungs, the connective tissue was removed, and arteries were opened for endothelial cell removal with a cotton swab in cold (4°C) HBSS. The arteries were allowed to recover for 30 min in cold HBSS followed by 20 min in reduced-Ca^2+^ HBSS (20 μM CaCl_2_) at room temperature. The tissue was enzymatically digested for 10-20 minutes at 37°C in reduced-Ca^2^ HBSS containing collagenase (type I; 1750 U/ml), papain (9.5 U/ml), bovine serum albumin (2 mg/ml) and dithiothreitol (1 mM). After digestion, the tissue was pipetted up and down in Ca^2+^-free HBSS and PASMCs were plated in growth media consisting of SmCM complete media (ScienCell) supplemented with the Lonza Smooth Muscle Cell bullet kit (cc-4149), and 1% penicillin/streptomycin. All cells were used at passage 1-2 and were placed in basal media, consisting of SmCM complete media (ScienCell) supplemented with 0.3% FBS and 1% penicillin/streptomycin for 24-48 hr before beginning experiments. Assessment of smooth muscle culture purity was performed using intracellular calcium concentration ([Ca^2+^]_i_) responses to 80 mM KCl and immunofluorescence in cells stained with smooth muscle-specific α-actin (SMA; 1:400, #19245s, Cell signaling) or smooth muscle myosin heavy chain (SMMHC; 1:100, ab125884, Abcam) and DAPI (nuclear stain; 1:10,000 in PBS, Invitrogen). Only cultures where >90% of cells exhibited at least 50 nM increase in [Ca^2+^]_i_ in response to KCl and/or positive SMA or SMMHC expression were used for these studies.

### Hydrogen peroxide (H_2_O_2_) exposures

At approximately 60% confluence, PASMCs were placed into basal media for 24 hr (Nor or SuHx) or 48 hr (post adenoviral infection to ensure proper protein overexpression). Cells were washed 3 times with PBS before being placed in serum-free SmCM media containing freshly made H_2_O_2_ (250 μM) and incubated for 24 hr before additional measurements (apoptosis, caspase activity or immunoblot).

### Hoechst staining

PASMCs were stained with Hoechst 33342 dye (1:5000; H3570; Invitrogen) for 15-30 min at 37° C to visualize apoptotic cells. Apoptosis was identified by observing alterations in chromatin morphology (i.e., condensation) upon staining. For each sample, 9-12 images of randomly chosen fields were captured at 20X via fluorescent microscopy (Olympus IX51) by an investigator blinded to treatment conditions and analyzed using ImageJ. Cells with normal and condensed chromatin were automatically identified and counted by DenoiSeg (https://github.com/juglab/denoiseg/), an open-source machine learning image segmentation algorithm. A custom model was developed using manually labeled training images in FIJI using an Nvidia Quadro RTX 4000 GPU using TensorFlow 1.15, CUDA 10.0, and cudnn 7.6.5 as previously described^37^. Apoptosis was determined as the percent of apoptotic cells over total cells and counts for each image within a sample were averaged to obtain a single value.

### Caspase activity assay

Caspase 3/7 activity was measured using the Caspase-Glo^®^ 3/7 Assay System (G8090, Promega) per the manufacturer’s instructions. Briefly, equal amounts of cell lysate (5 µg) diluted in T-PER™ were placed into the wells of a 96 well plate in duplicates for each sample. 50 μl of Proluminescent caspase-3/7 DEVD-aminoluciferin substrate was added to each well and read with a luminometer at 5 min intervals up to 60 min. Luminescence values within the linear portion of the curve (typically at 10 min) for each sample were averaged to obtain a single value, background subtracted and normalized to control within the same plate.

### Immunoblotting

Total protein was extracted from PASMCs in ice-cold T-PER buffer (ThermoFisher, #78510) containing cOmplete protease inhibitors (Roche Diagnostics) and phosphatase inhibitors cocktails (Sigma). Quantification of proteins was performed using BCA protein assay (Pierce) and equal amounts of total protein per lane were separated by electrophoresis through an 8% or 10% SDS-PAGE gel and transferred onto polyvinylidene difluoride membranes. Membranes were incubated in 5% non-fat dry milk in Tris-buffered saline containing 0.2% Tween 20 to block nonspecific binding sites before being probed with primary antibody against NHE1 (1:1000, Millipore Sigma-Aldrich, #MAB3140, specificity for NHE1 validated via overexpression and knockdown), glucose transporter type 1 (GLUT1;1:1000, ab115730, specificity confirmed in GLUT1 knockdown^38^), phosphorylated-p38 (1:1000, cell signaling, #9215) and p38 (1:1000, Cell Signaling, #9212 confirmed specificity by knockdown^39^), Na^+^/K^+^ ATPase (1: 1000, Millipore Sigma #05-369, confirmed specificity by knockdown^40^). Bound antibody was probed with horseradish peroxidase-conjugated anti-rabbit or anti-mouse secondary antibody (1:10,000; 52200336 and 52200341, SeraCare Life Sciences) and detected by enhanced chemiluminescence (Clarity Western ECL Substrate, 170-5061) using a Chemidoc gel imaging system (BioRad). Membranes were then stripped and re-probed for β-tubulin (1:10,000, T7816, Sigma) as a housekeeping protein. Protein levels were quantified by densitometry using ImageJ. All antibodies were validated before purchase or by using appropriate negative controls.

### Surface biotinylation

Cell surface biotin-labeling reagent (0.5 mg/ml, EZ-Link Sulfo-NHS-LC-Biotin, Pierce) was added to cultured PASMCs for 1 hr and quenched by addition of 50 mM Tris (pH 8.0). Cells were then washed with PBS and lysed with solubilizing buffer (20 mM Tris HCI at pH 8.0, 200 mM NaCl, 1 % Triton x-100, 1 % Sodium Deoxycholate, 5 mM EDTA) containing protease inhibitors and total sample protein concentration determined via BCA protein assay. Equal amounts of protein were incubated with streptavidin beads overnight (Streptavidin Agarose Resins; Thermo Scientific) to immunoprecipitate biotinylated proteins. Samples were washed four times in solubilization buffer and salt-free buffer (20 mM Tris HCl PH 8.0, 1 % Triton x-100, 5 mM EDTA) to facilitate removal of non-specific proteins. Biotinylated protein was eluted in Laemmli sample buffer (BioRad) containing 3% SDS and 10% β-mercaptoethanol (BME) for 30 min at 37°C, subjected to immunoblotting as described above.

### Generation of adenoviral constructs

Generation of adenoviral constructs containing a hemagglutinin (HA)-tagged wild-type NHE1 (AdNHE1-WT) or NHE1 with a mutation in the ezrin binding (AdNHE1-EM) was previously described^25,29,41^. For the ion translocation mutant (AdNHE1-E262I), we cloned a cDNA for human NHE1 into pCMV-Sport6 (obtained from the MGC Collection-Genbank Accession #<u>BC051431</u>) through Open Biosystems, Inc. (Birmingham, ALA). A small fragment of the cDNA encoding nucleotides 1638–2582 was generated by PCR and an epitope tag (2 x HA) was added to the C-terminus of the protein. DNA sequence was confirmed, and the extraneous 3′ untranslated region, 5′ UTR sequences and the remainder of the coding sequence (nucleotides 120–1637) was replaced by a PCR product into the KpnI/SalI sites of pCMVSport 6.

The NHE1 cDNA was transferred by recombination cloning into pDONR221 and then into pAdDEST-V5 to generate the plasmid, pAdCMV-NHE1-2XHA (AdNHE1-WT) using manufacturer’s instructions (InVitrogen). Site directed mutagenesis was performed to the NHE1 construct where a single amino acid substitution at position 262 was made from a glutamate (E) into isoleucine (I) as previously described^29,41,42^ for the generation of AdNHE1-E262I. Transfection into 293A cells and isolation of cell extracts showing cytopathic effects allowed for the production of replication defective virus. This cell extract was then used to amplify the virus. Viral titer was determined using the Adeno-X™ Rapid Titer Kit (Clontech) and virus purification was performed using Adenopure Filter system (PureSyn, Inc.). At approximately 60% confluency, PASMCs grown in culture dishes were placed in basal media supplemented with 0.3% FBS and 1% penicillin/streptomycin and infected with 50 Multiplicity of Infection (MOI) of a control adenovirus containing GFP (AdGFP), AdNHE1-WT (HA-tagged wild-type), AdNHE1-E262I (HA-tagged NHE1 with a mutation in the ion translocation site) or AdNHE1-EM (HA-tagged NHE1 with a mutation in the ezrin binding) for 48 hr at 37°C before experiments.

### NHE activity

NHE activity in PASMCs was measured using the ammonium pulse technique as previously described^7,21–23,43,44^. PASMCs were plated onto a coverslip and incubated with the cell permeant pH sensitive dye 2’,7’-bis(carboxyethyl)-5(6)-carboxyflourescein (BCECF-AM) for 1 hr prior to experiments. The cells were placed in a perfusion chamber perfused with HEPES-buffered physiologic salt solution (PSS) containing (in mM): 130 NaCl, 5 KCl, 1 MgCl_2_, 1.5 CaCl_2_, 10 glucose, 20 HEPES-Tris, with pH adjusted to 7.4 using NaOH. PASMCs were excited with light filtered at 490 and 440 nm and fluorescence emitted from the PASMCs was detected at 530 nm. The ratio of 490 to 440 nm emission was calculated, and pH_i_ was estimated from *in situ* calibration after each experiment. For calibration, PASMCs were perfused with a solution containing (in mM): 106 KCl, 1 MgCl_2_, 1.5 CaCl_2_, 10 glucose, 20 HEPES-Tris and 0.01% nigericin to allow pH_i_ to equilibrate to external pH. A two-point calibration was created based on the fluorescence values measured using solutions with pH of 6.5 and 7.5 (adjusted with KOH). Intracellular H^+^ ion concentration ([H^+^]_i_) was determined from pH_i_ using the formula: pH_i_ = -log ([H^+^]_i_). Data was collected from 10-30 cells per coverslip and averaged to obtain a single value for each biological replicate per experiment

### Thioflavin T (ThT)

PASMCs were placed in basal media 24 hr prior to the experiment. Cells were then washed with PBS and placed in serum-free media and treated with either vehicle (PBS) or 250 μM of H_2_O_2_ to stimulate ER stress. Following treatment for 24 hr, PASMCs were incubated with Thioflavin T (ThT, 5 μM) or vehicle (PBS) for 5 minutes and fluorescence measured at 20X via fluorescent microscopy (Olympus IX51). At least 4 images per group were obtained under excitation 490, emission 520, by an investigator blinded to treatments/conditions. Fluorescence was measured by mean gray value measurements on ImageJ of at least 9 cells per image, background subtracted and averaged to obtain a single value.

### siRNA

Depletion of endogenous NHE1 was achieved using siRNA specifically targeting NHE1 (siNHE1; Horizon Discovery) and nontargeting (siNT; control) siRNA obtained as a “smart pool” (Horizon Discovery, D-001810-10-50). PASMCs were incubated with 100 nM of siRNA for 16 hr in serum-and antibiotic-free media, after which serum was added to the media for a total concentration of 0.3% FBS. PASMCs were incubated under these conditions for 8 hr, and then media was replaced. Cells were incubated for an additional 24 hr in basal media (0.3% FBS) prior to experiments. NHE1 knockdown was confirmed by immunoblotting.

### Data Analysis

Data are expressed as scatter plots with bars representing mean ± SD. Each dot represents a separate experimental run, and since all experimental runs were performed on tissue/cells from different animals, “n” also refers to the number of animals. All data were tested for normality and equal variance prior to running statistical tests. Data that were not normally distributed were log (10) transformed and retested for normality and equal variance prior to running statistics. Statistical comparisons were performed using Students *t*-test for data in two groups, or one-or two-way ANOVA with a Holm-Sidak post hoc test for multiple group comparisons. For all imaging experiments (immunofluorescence, Hoechst staining), images across groups were collected at the same time, using the same conditions within an experiment and the investigator obtaining the images and performing counting was blinded to treatments/groups.

## Results

### Increased ER stress and suppressed apoptotic pathway in SuHx PASMCs

Measurements of RV/LV+S weight in the SuHx rats was higher than in Nor controls confirming the development of right ventricle hypertrophy and PH in SuHx animals (Fig 1A, Supplemental Table 1). We also confirmed PASMCs from SuHx rats had higher NHE activity compared to Nor controls (Fig 1B) measured via ammonium pulse technique, similar to data observed in mice with chronic-hypoxia induced PH^22–24^. Immunoblotting of surface proteins following surface biotylation assay showed a small but significant increase in NHE1 expression normalized to Na^+^/K^+^ ATPase in PASMCs from SuHx rats when compared to Nor PASMCs (Fig 1C). At baseline, Thioflavin (ThT) fluorescence was increased in PASMCs isolated from SuHx animals compared to Nor PASMCs, indicating elevated markers of ER stress in SuHx PASMCs (Fig 1D). Addition of H_2_O_2_ for 24 hr increased ThT in PASMCs isolated from Nor rats, while no further increase with H_2_O_2_ was observed in PASMCs from SuHx rats PASMCs from the SuHx animal model have been previously reported to be resistant to apoptosis^10^. To confirm these findings, we measured basal levels of apoptosis for both Nor and SuHx PASMCs measured via Hoechst staining (Fig 1E). Even with the increased levels of ER stress in PASMCs from SuHx animals at baseline, basal apoptosis was not statistically different in PASMCs between Nor and SuHx rats (Fig 1E). Following stimulation of apoptosis with H_2_O_2_ (250 µM) for 24 hr, apoptosis significantly increased in PASMCs from Nor animals. H_2_O_2_ induced apoptosis in PASMCs from SuHx animals but to a significantly lower extent than in Nor cells, confirming resistance to apoptosis even in the presence of exogenous oxidative stress.

**Fig 1.**
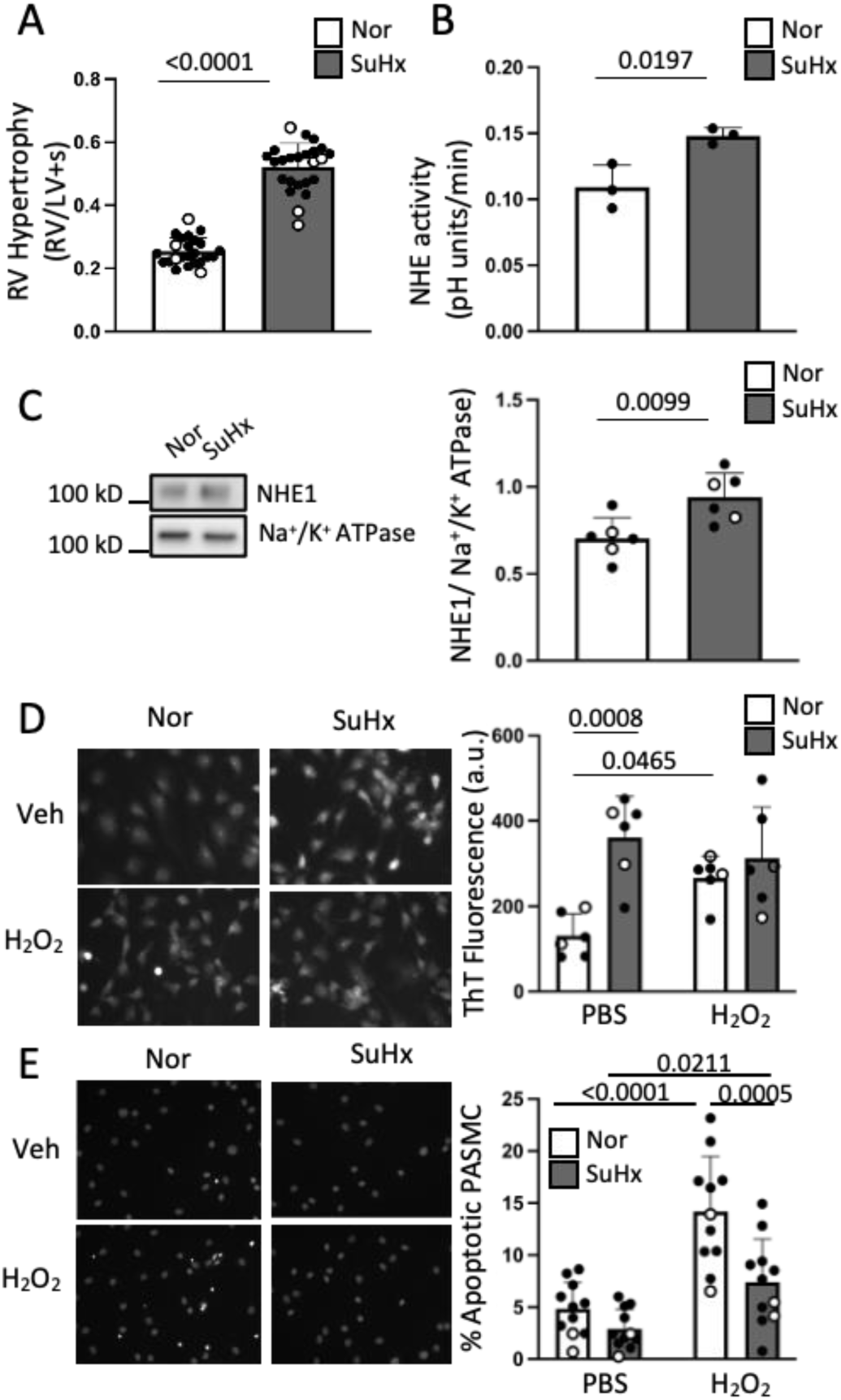
Hemodynamic and cellular changes in Sugen/ hypoxia (SuHx) rats. **A)** Bar and scatter plots show right ventricle over left ventricle + septum (RV/LV+S) weight ratio in normoxic (Nor) and SuHx rats (mean +/- SD, n=25) following the 5-week protocol. **B)** Bar and scatter plots show mean +/- SD for Na^+^/H^+^ exchanger (NHE) activity of pulmonary arterial smooth muscle cells (PASMCs) for Nor and SuHx rats (n=3). **C)** Representative immunoblot and bar and scatter plots graph show mean +/- SD of NHEĨ protein expression (n=5) measured in PASMCs of Nor and SuHx following surface biotinylation. Significance was assessed by unpaired two-tailed t-test for (A-C). **D)** Representative images and bar and scatter plots show mean +/- SD of Thioflavin T fluorescence (n=5) measured in PASMCs of Nor and SuHx in response to incubation with H_2_O_2_(25O µM; 24 hr) or vehicle (PBS). **E)** Representative images and bar and scatter plots show apoptosis (n=ll) measured via Hoechst staining in response to incubation with H_2_O_2_ (250 µM; 24 hr) or vehicle (PBS). Values are presented as percent of total cells. Significance assessed by two-way ANOVA with Holm-Sidak post hoc test. Interaction for **D)** p= 0.0176 and **E)** p= 0.0365. Black dots represent data from PASMCs from male rats, while white dots represent data from PASMCs from female rats.

### Role of NHE1 in PASMCs apoptosis

To explore the correlation between increased transmembrane NHE1, increased NHE activity and decreased apoptosis in PASMC from SuHx rats, we used 10 μM of ethyl-isopropyl amiloride (EIPA), a potent NHE inhibitor. Addition of EIPA significantly decreased NHE activity (Fig 2A, B) consistent with our previously reported results^7^.

**Fig 2.**
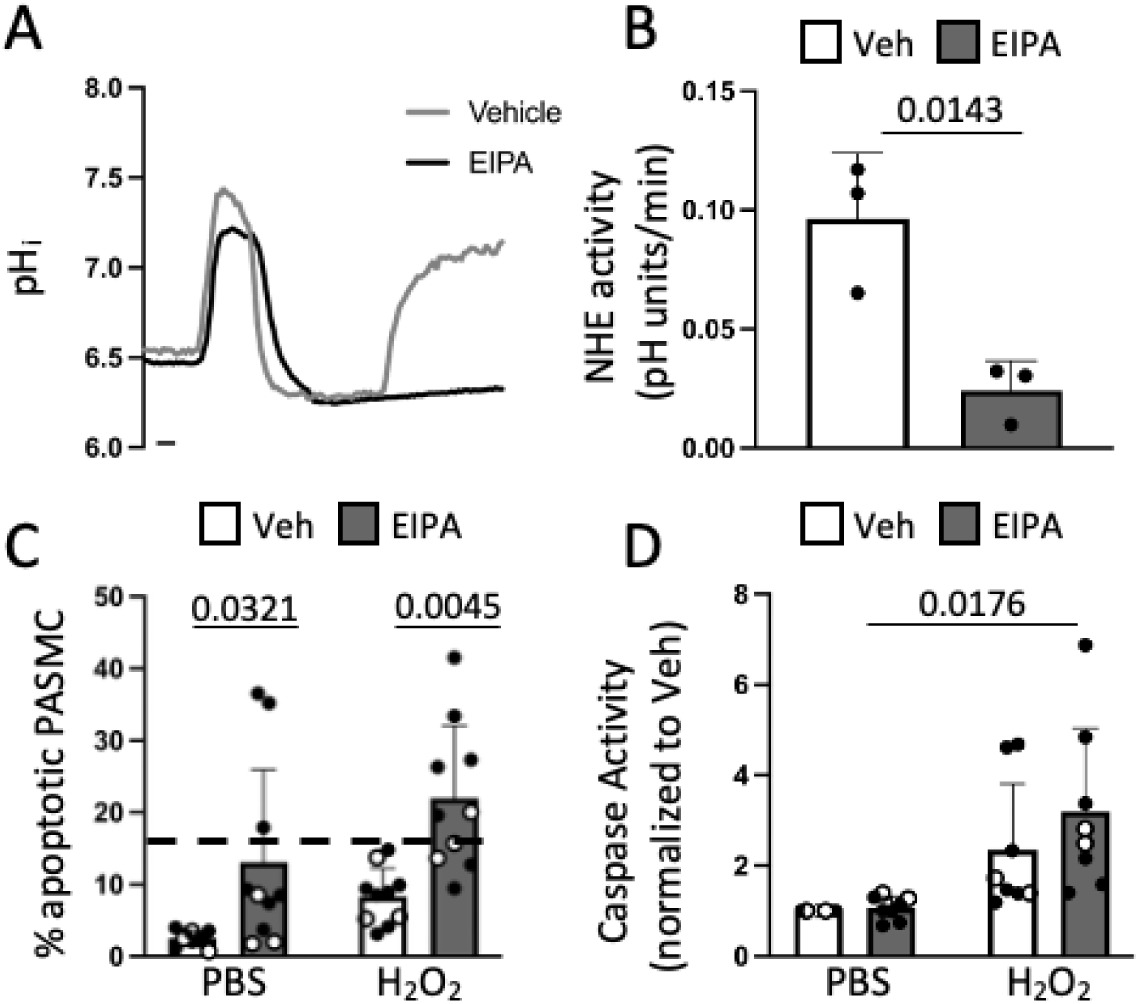
Effect of Na7H* exchanger (NHE) inhibition with ethyl isopropyl amiloride (EIPA) on resistance to apoptosis in Sugen/hypo×ia SuHx pulmonary arterial smooth muscle cell (PASMC). **A)** Representative traces and **B)** bar and scatter plots show NHE activity (mean +/- SD) in PASMCs following EIPA treatment (10 µM, 24 hr) or vehicle (DMSO) during the ammonium pulse protocol (n=3). Significance was assessed by unpaired two-tailed t-test. Bar and scatter plots show mean +/- SD for apoptosis in PASMCs from SuHx rats in response to stimulation with H_2_O_2_ (250 µM) or vehicle (PBS) following EIPA incubation (10 µM, 24 hr) measured by **C)** Hoechst staining (n=lO) and **D)** caspase activity (n=8). Values for Hoechst staining are presented as percent of total cells while values for caspase activity are normalized to PBS-treated SuHx cells. Significance assessed by two-way ANOVA with Holm-Sidak post hoc test. For Cļ, interaction p= 0.5663. For D), interaction p-0.3126. The dashed line represents the average value for normoxic control apoptosis percent following H_2_O_2_ treatment. Black dots represent data from PASMCs from male rats, while white dots represent data from PASMCs from female rats.

In SuHx PASMCs, pre-treatment with EIPA significantly increased apoptosis at baseline (Fig 2C), and in response to H_2_O_2_ (Fig 2D). We also measured PASMC caspase 3/7 activity with the Caspase-Glo^®^ 3/7 Assay System as a complementary measurement of apoptosis. At baseline, there was no change in caspase-induced luminescence between vehicle or EIPA treated groups. Compared to the PBS controls, H_2_O_2_ caused a significant increase in caspase activity in EIPA-treated, but not vehicle-treated, cells (Fig 2D).

Since EIPA is a non-isoform selective inhibitor of NHE, we repeated experiments with a NHE1-specific inhibitor, cariporide (10 μM). Similar to EIPA, cariporide significantly decreased NHE activity measured via the ammonium pulse experiment (Fig S1A, B). Surprisingly, following pre-treatment with cariporide, we observed no changes in apoptosis at either baseline or in response to H_2_O_2_ measured by Hoescht staining (Fig S1C) or Caspase-Glo^®^ 3/7 Assay System (Fig S1D).

Given the results with cariporide, we sought to confirm the specificity of the role of NHE1 in SuHx PASMC apoptosis using siRNA to knockdown NHE1 protein expression. NHE1 protein expression (Fig 3A,B) and NHE activity (Fig 3C) were significantly decreased following siNHE1 treatment when compared to siNT. Silencing NHE1 increased apoptosis in PASMCs from SuHx rats at baseline when compared to siNT (Fig 3D). Treatment with 250 μM of H_2_O_2_, significantly increased apoptosis in the siNHE1 treated PASMCs when compared to the siNT (Fig 3D). While caspase 3/7 activity in response to H_2_O_2_ was not different in the siNT control groups, H_2_O_2_ stimulation increased caspase 3/7 activity in the siNHE1 group (Fig 3E).

**Fig 3.**
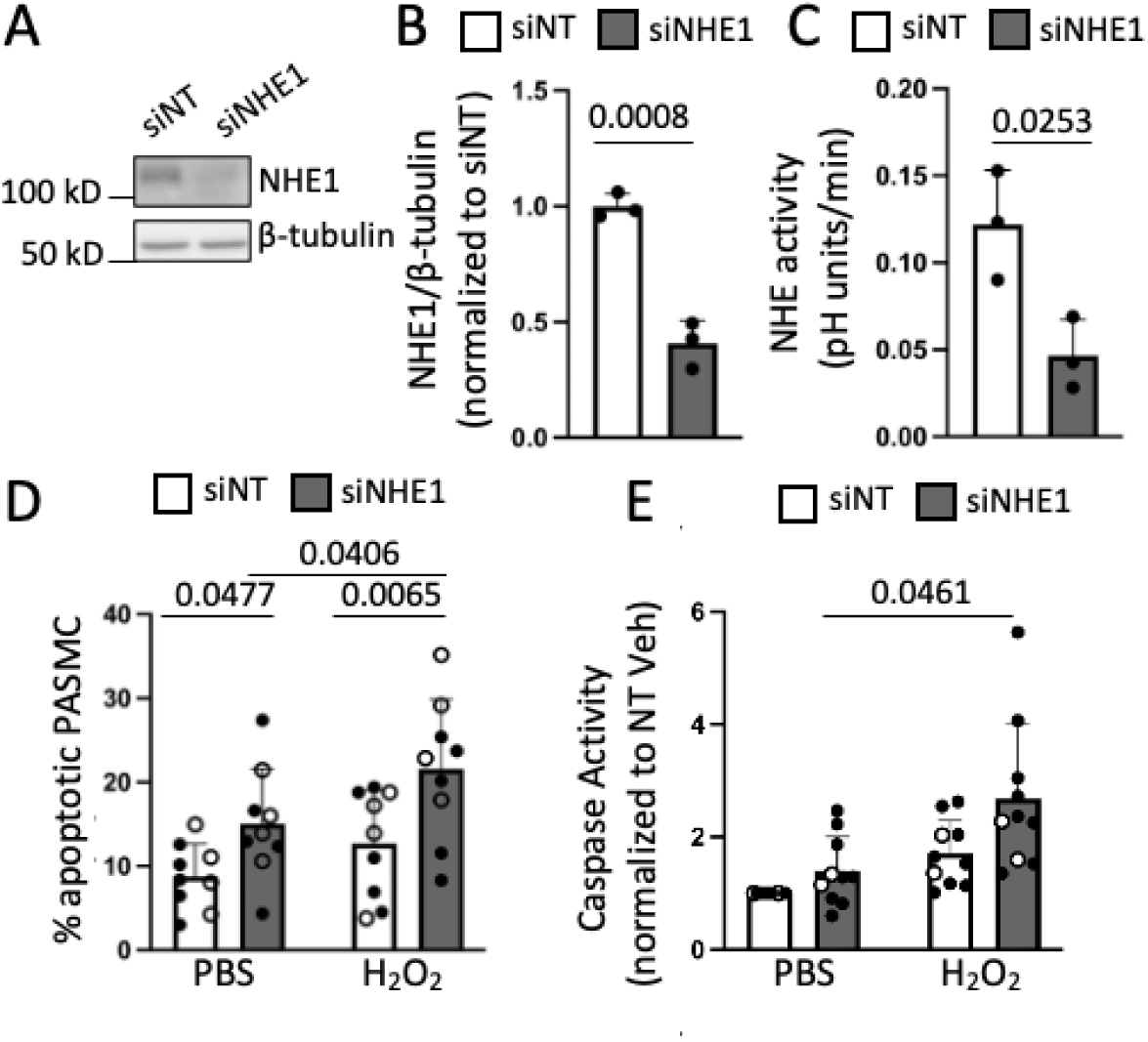
Effect of Na^+^/H^+^ exchanger 1 (NHE1) depletion on apoptosis in Sugen/hypoxia (SuHx) pulmonary arterial smooth muscle cells (PASMCs). **A)** Representative blots show NHE1 protein levels in PASMCs compared to cells transfected with siRNA targeted to NHE1 (siNHEl) or non-targeting siRNA (siNT). **B)** Bar and scatter plots represent mean + SD values normalized to siNT (n = 3) for NHE1 protein levels presented as NHEl/β-tubulin ratio. Significance was assessed via unpaired two tailed ř-test. **C)** Bar and scatter plots show NHE activity (mean +/- SD) for siNHEl- and siNT-treated cells (n=3). Significance assessed by unpaired two tailed t-test. **D)** Bar and scatter plots show mean +/- SD for apoptosis in rat PASMCs from SuHx animals (n=9) following siNHEl or siNT treatment and stimulation with H_2_O_2_ (250 µM, 24 hr) or PBS measured by Hoechst staining and **E)** caspase activity (n=lO). Values for Hoechst staining are presented as percent of total cells while values for caspase activity are normalized to siNT PBS SuHx cells. Significance assessed by two-way ANOVA with Holm-Sidak post hoc test. For **D)**, interaction p = 0.9382. For **E)**, interaction p = 0.4982. Black dots represent data from PASMCs from male rats, while white dots represent data from PASMCs from female rats.

### Exploring the role of p38 in apoptosis resistance of PH-PASMCs

To elucidate the mechanism underlying NHE1-mediated apoptosis resistance in PASMCs, we analyzed the p38 MAPK pathway. The p38 MAPK pathway is known to be activated by reactive oxygen species in various cell types, promoting apoptosis^45,46^. Moreover, because activation of p38 has been reported to be pH-dependent in cancer cells^47^, we assessed the relationship between NHE1 inhibition and p38 activation. As expected, H_2_O_2_ increased phosphorylation of p38 (p-p38) in Nor PASMCs. However, H_2_O_2_ had no effect on p-p38 in PASMCs from SuHx rats (Fig 4A). In PASMCs from SuHx rats, incubation with EIPA or silencing NHE1 restored phosphorylation of p38 in response to H_2_O_2_ (Fig 4B, C).

**Figure 4.**
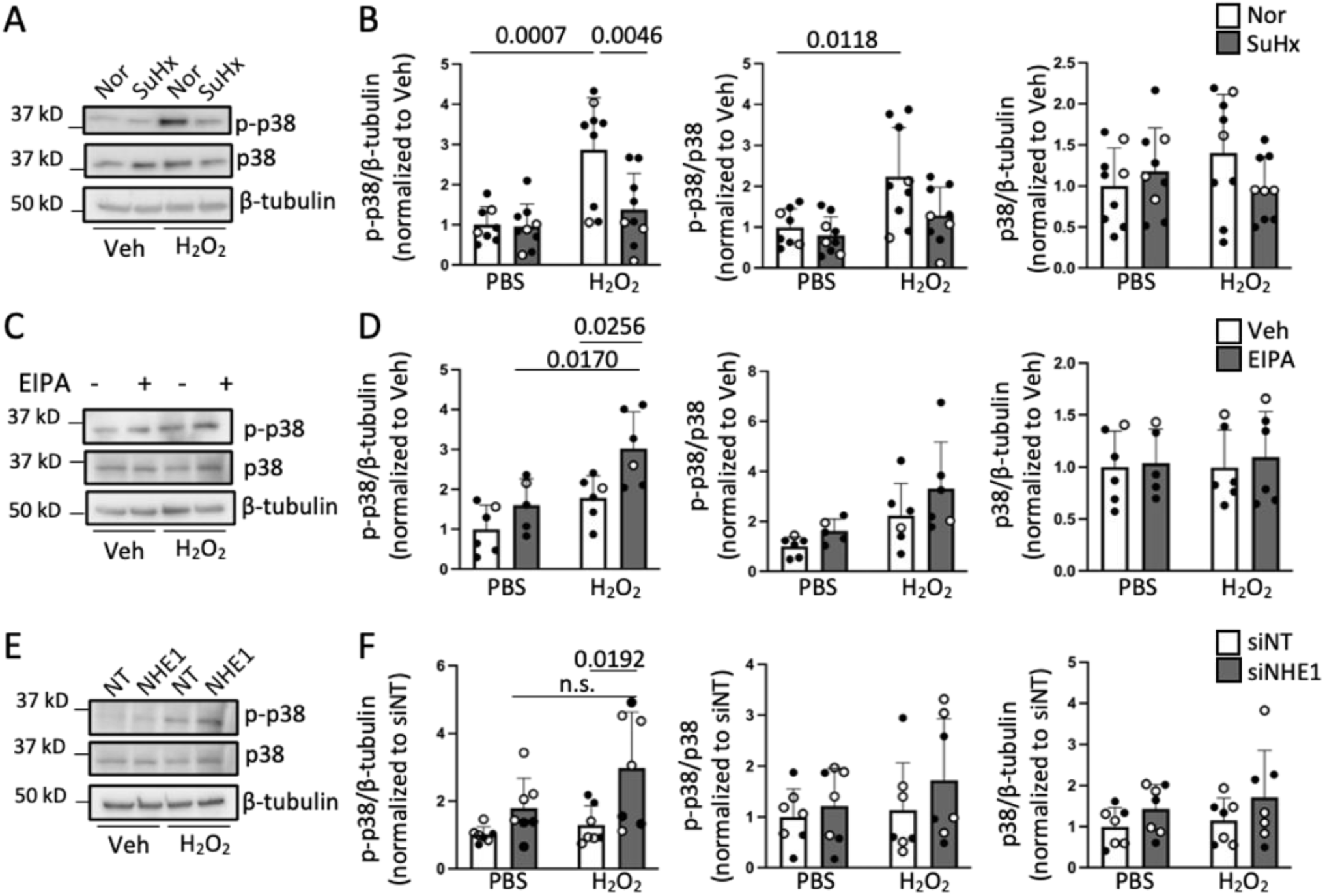
Phosphorylation of p38 following apoptotic stimuli in control cells and Sugen-Hypoxia (SuHx) pulmonary arterial smooth muscle cells (PASMCs) **A)** Representative immunoblots and **B)** quantification bar and scatter plots show mean + SD values (n = 9) for phosphorylated p38 (p-p38) protein expression following H_2_O_2_ (250 µM, 24 hr) or vehicle (PBS) treatment in normoxic (Nor) and SuHx PASMCs. Protein levels are presented as ratio of p-p38/β-tubulin, p-p38/p38 and p38/β-tubulin. **C)** Representative immunoblots and **D)** quantification bar and plots represent mean ±SD values (n = 5-6) for protein levels presented as ratio of p-p38/β-tubulin, p-p38/p38 and p38/β-tubulin following treatment with 10 µM EIPA or vehicle (DMSO) and apoptotic stimulation with H_2_O_2_ (250 µM, 24 hr) or vehicle (PBS). **E)** Representative immunoblots and **F)** quantification bar and plots represent mean + SD values (n = 7) for protein levels presented as ratio of p-p38/β-tubulin, p-p38/p38 and p38/β-tubulin following NHE1 silencing with siRNA targeted to NHE1 (siNHEl) or non-targeting control (siNT) and apoptotic stimulation with H_2_O_2_ (250 µM, 24 hr) or vehicle (PBS). Significance assessed by by two-way ANOVA with Hollm-Sidak post hoc test. Interaction for **B)** p = 0.0218, p=0.1611 and p=0.0992, for (D) p=0.2952, p=0.6637 and p=0.8530, for **F)** p=0.2405, p=0.5885 and p=0.8122. Black dots represent data from PASMCs from male rats, while white dots represent data from PASMCs from female rats.

To determine whether increased phosphorylation of p38 in response to EIPA mediated the EIPA-induced increase in susceptibility to apoptosis in PASMCs from SuHx rats, we inhibited the activity of p-p38 using 10 μM of SB203580 and measured apoptosis in EIPA-treated SuHx cells. Treatment of SB203580 prevented the increase in SuHx apoptosis induced by EIPA when measured by Hoechst staining (Fig 5A). Unexpectedly, we did not observe any changes in caspase activity following 24 hr of apoptotic stimulation (Fig 5B).

**Fig 5.**
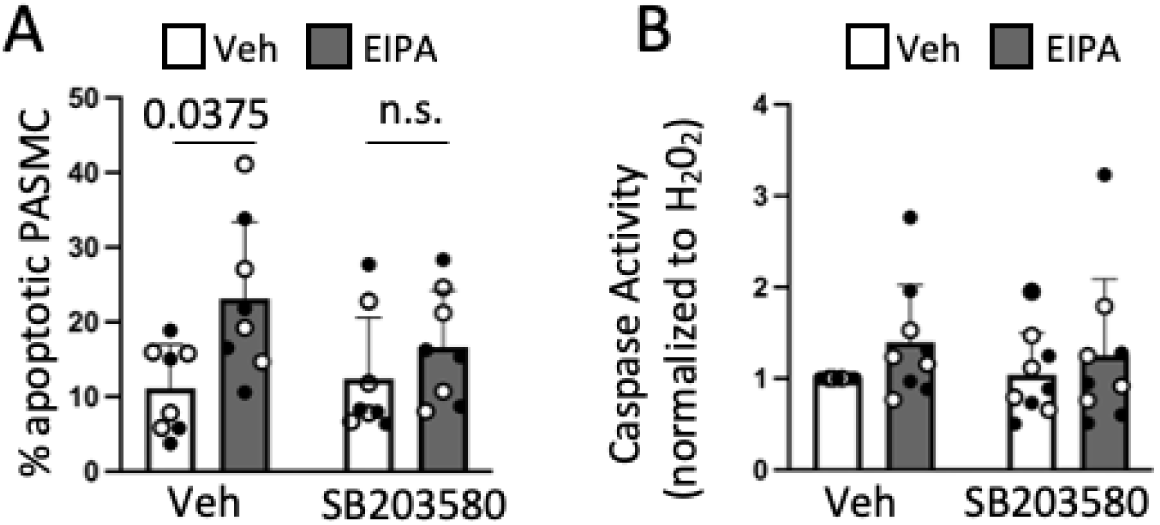
Effect of inhibition phosphorylated p38 activity with SB203580 in Sugen/hypoxia (SuHx) pulmonary arterial smooth muscle cell (PASMC) apoptosis following H_2_O_2_ stimulation. Bar and scatter plots show mean +/- SD (n=8) for apoptosis in rat PASMCs from SuHx animals in response to stimulation with H_2_O_2_(2 50 µM, 24 hr) following incubation with EIPA (10 µM, 24 hr) or vehicle (DMSO) and SB203580 (10 µM, 24 hr) or vehicle (DMSO) measured by **A)** Hoechst staining and **B)** caspase activity. Values for Hoechst staining are presented as percent of total cells while values for caspase activity are normalized to SuHx vehicle cells. Significance assessed by two-way ANOVA with Holm-Sidak post hoc test. For A), interaction p= 0.1873 For B), interaction p= 0.7157. Black dots represent data from PASMCs from male rats, while white dots represent data from PASMCs from female rats.

### Effect of augmented NHE1 expression on PASMC apoptosis

Given the role of NHE1 in apoptosis resistance in PASMCs from SuHx rats, we next tested whether increasing levels of NHE1 was sufficient to induce apoptosis resistance in PASMCs and, if so, through which mechanism. We increased NHE1 expression using AdNHE1-WT, AdNHE1-E262I (ion translocation mutant) or AdNHE1-EM (ezrin binding mutant) with AdGFP as control for infection.

Protein level and localization of expressed NHE1 were assessed by surface biotinylation where all 3 NHE1 constructs were expressed at similar levels that were significantly higher than AdGFP (Fig 6A, B). NHE activity (Fig 6C) was increased in PASMCs expressing AdNHE1-WT and AdNHE1-EM, but not AdNHE1-E262I, confirming functionality of AdNHE1-WT and AdNHE1-EM and the inability of AdNHE1-E262I to translocate ions. Infection with AdNHE1-WT inhibited H_2_O_2_– induced apoptosis, measured by Hoechst staining and caspase 3 activity, compared to cells infected with AdGFP (Fig 6D, E). Surprisingly, AdNHE1-E262I and AdNHE1-EM also inhibited PASMC apoptosis following H_2_O_2_ stimulation (Fig 6D, E). These findings suggest increasing NHE1 is sufficient to suppress PASMC apoptosis and that the mechanism is independent of pH_i_ regulation or interaction with ERM proteins. Moreover, consistent with the association between increased p38 phosphorylation and apoptosis, increasing NHE1 expression with AdNHE1-WT, AdNHE1-EM or AdNHE1-E262I in control PASMCs also significantly reduced the phosphorylation of p38 in H_2_O_2_-stimulated PASMCs when compared to AdGFP control (Fig 7).

**Fig 6.**
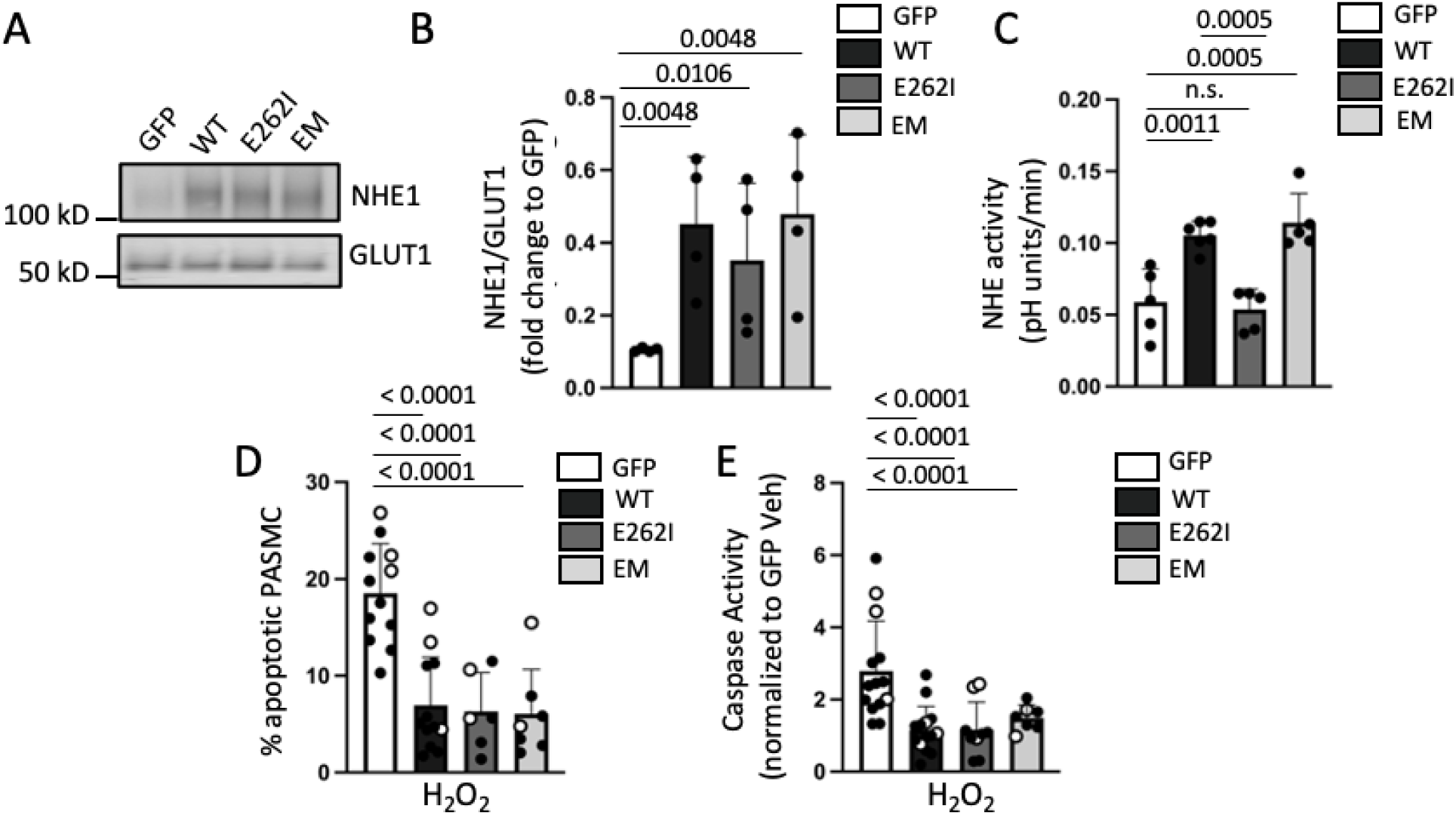
Apoptosis levels in pulmonary arterial smooth muscle cells (PASMCs) from control rats following overe×pression of wild-type Na*/H^+^ exchanger 1 (WT), ion-translocation mutant (E262l) and ERM binding mutant (EM). **A)** Representative immuπoblots show NHE1 protein levels post adenovirus infection following surface biotinylation. **B)** Bar and scatter plots represent mean ±SD values (n=4 per group) for NHE1 protein levels presented as NHE1/GLUT1 ratio. Data was tested for normality and log transformed before running one-way ANOVA with Holm-Sidak post hoc test. **C)** Bar and scatter plots show (mean +/- SD) for NHE activity of PASMC following adenovirus infection. Significance assessed by one-way ANOVA with Holm-Sidak post hoc test. **D)** Bar and scatter plots represent mean±SD for apoptosis measured via Hoechst staining (n=5-12) and **E)** via caspase activity (mean +/- SD) (n=7-14) in rat PASMCs from control animals following adenovirus infection in response to H_2_O_2_ (250 µM, 24 h). Values for apoptosis are presented as percent of total cells while values for caspase activity are normalized to AdGFP Veh cells (data not shown). Significance assessed by one-way ANOVA with Holm-Sidak post hoc test. Black dots represent data from PASMCs from male rats, while white dots represent data from PASMCs from female rats.

**Figure 7.**
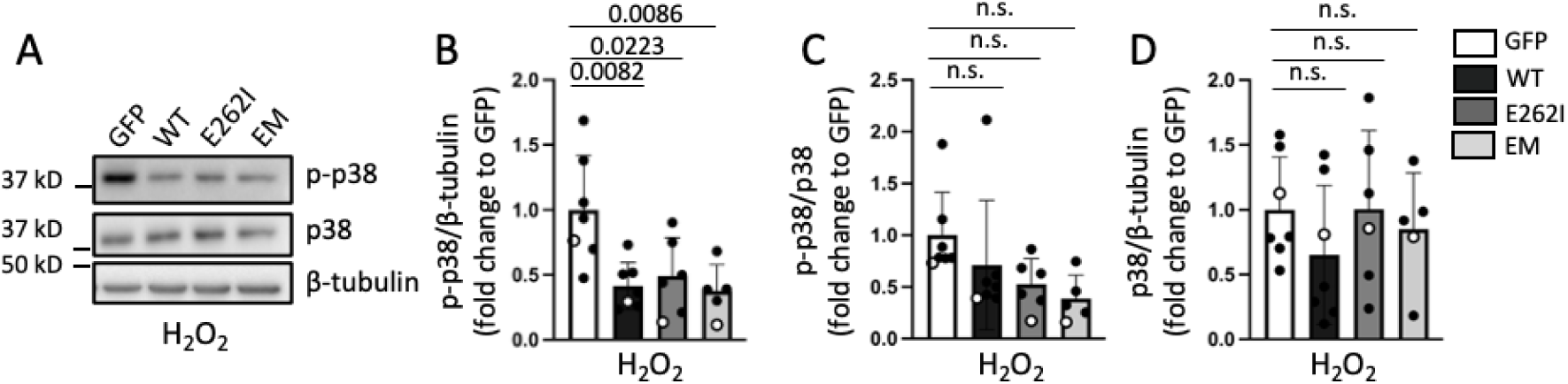
Effects of overexpression of WT and mutant Na‘/H* exchanger 1 (NHE1) constructs on H_2_O_2_-induced phosphorylation of p38 (p-p38) in control pulmonary arterial smooth muscle cells (PASMCs). **A)** Representative immunoblots show phosphorylation of p38 following H_2_O_2_(250 µM, 24 hr) treatment in control cells following overexpression of GFP control and NHE1 constructs (WT, E262l and EM). Quantification bar and scatter plots show phosphorylation of p38 presented as mean ±SD ratio of **B)** p-p38/β-tubulin, **C)** p-p38/p38 and **D)** p38/β-tubulin following virus infection and H_2_O_2_ treatment (n=5-7). Significance assessed by one-way ANOVA with Holm-Sidak post hoc test. For **B)** p=0.0002, for **C)** p = 0.2796, for **D)** p = 0.3065. Black dots represent data from PASMCs from male rats, while white dots represent data from PASMCs from female rats.

## Discussion

In this study, we describe the role of NHE1 in mediating changes in PASMC apoptosis from control and SuHx rats. We confirmed findings of previous studies demonstrating upregulation of NHE activity^7,44^ and resistance to apoptosis^10^ in SuHx PASMC. The results from this current study indicate that NHE1 is necessary for resistance to apoptosis in PASMCs from SuHx rats and that increased NHE1 expression is sufficient to confer apoptosis resistance in control PASMCs. Additionally, these results suggest neither pH_i_ regulation nor interaction between NHE1 and ezrin are essential for NHE1-induced apoptosis resistance. Finally, consistent results in SuHx and control PASMCs indicate the mechanism underlying NHE1-induced resistance to apoptosis involves p38 phosphorylation.

Increased NHE activity has been reported in PASMC from different models of PH^7,23,24,44^. PASMCs from CH animals have increased NHE activity as well as upregulation of NHE1 at the protein and mRNA levels^23^. In PASMCs from SuHx rats, our lab previously reported an increase in NHE activity, but surprisingly, in the absence of an increase in NHE1 total expression^7^. In this study, we confirmed higher NHE activity in SuHx PASMCs and increased NHE1 protein expression in the plasma membrane, likely explaining increased NHE activity in SuHx PASMCs. The mechanism through which PASMCs from SuHx rats can upregulate NHE1 expression at the plasma membrane level independent of changes in the total cell content is still unclear. One possibility is that the pathway controlling NHE1 turnover at the plasma membrane may be altered. Recent findings show NHE1 undergoes endocytosis via ubiquitylation facilitated by the adapter protein β-arrestin-1^48^. β-arrestin-1 expression is reduced in lung lysates from patients with PAH^49^, leading us to speculate increased surface NHE1 may result from reduced NHE1 ubiquitination and endocytosis in SuHx PASMCs. Further experiments are needed to validate this hypothesis.

In PASMCs from Nor rats, increasing ER stress by application of H_2_O_2_ caused apoptosis as expected^50^. Interestingly, in PASMCs from SuHx rats, ER stress was increased at baseline, consistent with findings in other models of PH^19,20,51–53^. Despite increased ER stress, a condition that should produce endogenous pro-apoptotic stimuli, apoptosis at baseline was not elevated in PASMCs from SuHx rats, suggesting SuHx PASMCs adapted to increased ER stress and/or are not capable of responding to endogenous apoptotic stimuli. Similarly, H_2_O_2_ treatment failed to further increase ThT levels in PASMCs from SuHx rats, indicating that ER stress was already maximally elevated in these cells.

Because of the important role of NHE1 in migration and proliferation of SuHx^7^ PASMC and in apoptosis in cancer models^54^, we tested whether increased NHE1 surface expression or increased NHE activity in PASMCs from SuHx rats regulated apoptosis susceptibility. Previous data showed EIPA inhibits NHE activity in rat PASMC^7^, which we confirmed. EIPA reversed resistance to apoptosis in PASMC from SuHx animals at both baseline and following stimulation with H_2_O_2_, indicating NHE is required for apoptosis resistance in SuHx PASMCs. These results suggest NHE inhibition with EIPA sensitized PASMCs from SuHx rats to the high levels of endogenous ER stress at baseline, allowing apoptosis to proceed. One limitation from this experiment is that EIPA inhibits multiple NHE isoforms, including NHE1, NHE2, NHE3, and NHE5 with varying affinities^55^. Publicly available single-cell transcriptomics data sets (proteinatlas.org) reveal lung smooth muscle cells express NHE1, NHE5, NHE6, NHE8 and NHE9^56^. The NHE6, NHE8, and NHE9 isoforms are mainly associated with intracellular membranes such as endosomes, the trans-Golgi network, and secretory vesicles^57,58^. NHE5 localizes to both recycling endosomes and the plasma membrane^58^, and while transcript expression is very low in lung SMCs, we could not rule out the possibility that EIPA could be exerting actions inhibiting NHE5. Thus, to confirm the role of NHE1 in SuHx PASMC resistance to apoptosis, we used siRNA targeting NHE1 to decrease NHE1 protein expression and NHE activity. Even with incomplete knockdown, siNHE1 increased apoptosis similar to our results with EIPA. We also noted a small increase of about 5.5% in apoptosis at baseline in the siNT-treated group compared to untreated cells, which speculate might be due to ER stress induced by siNT treatment, as noted in other cells^59,60^.

Given our findings that inhibiting NHE activity or NHE1 silencing increased apoptosis in SuHx PASMCs, we next tested whether elevated NHE1 expression was sufficient to induce apoptosis resistance in control PASMCs. Expressed wildtype NHE1 protein properly translocated into the plasma membrane and increased NHE activity, and was able to prevent H_2_O_2_-induced apoptosis, confirming a role for NHE1 in regulating PASMC apoptosis and suggesting an elevation in NHE1 levels alone is sufficient to induce PASMC resistance to apoptosis.

It is well known cellular acidification is an important early event in apoptosis that facilitates caspase activation and optimal activity^61–63^. Taken together with our EIPA data, we hypothesized that higher amounts of NHE1 prevents apoptosis via increased NHE activity and inhibition of the initial cellular acidification. An NHE1 mutant (AdNHE1-E262I) that lacks ion translocation capacities^29,41,42^ but does not affect the ability of NHE1 to interact with the actin cytoskeleton^41^ properly translocated to the plasma membrane but did not increase NHE activity. Surprisingly, this mutant also rendered the PASMCs resistant to apoptosis. These results suggest the mechanism by which NHE1 mediates PASMC apoptosis is independent of pH_i_ regulation and may explain why inhibiting NHE1 activity with cariporide did not affect PASMC apoptosis. While both EIPA and cariporide are well used drugs known inhibit NHE ion transport, our results suggest that their mechanism of action is distinctive and EIPA may have an effect on NHE1 independent of exchanger activity. While studies on their putative binding sites exist^64,65^, to our knowledge, none of them explore confirmational changes that could happen in the cytosolic tail region of NHE1 following pharmacological inhibition, which could impact other non-canonical roles of NHE1, protein-protein interactions or different signaling pathways.

In addition to its important role in pH_i_ regulation via the transport of ions, NHE1 is involved in many other cellular pathways via non-canonical interaction in its C-terminus cytosolic tail domain^29,41^. Our lab previously published an essential role of NHE1 in PASMC migration and proliferation via protein-protein interactions between NHE1 and phosphorylated ezrin^25^. In apoptosis, the NHE1-ezrin interaction has been shown to be protective against apoptosis in other cell types^30,31^. We found in PASMCs that mutating the ERM motif in the cytosolic C-terminus of NHE1 disrupted NHE1-ezrin-actin binding^25^ but maintains the ability of NHE1 to translocate Na^+^ and H^+^ ions^41^. Similar to wildtype NHE1, overexpression of the AdNHE1-EM mutant increased NHE activity and resistance to apoptosis in PASMC, suggesting NHE1-induced resistance to apoptosis in PASMC is independent of the NHE1-ezrin protein interaction. These experiments demonstrate the mechanism of action of NHE1 in PASMC apoptosis does not require two of the most well-known roles of NHE1.

Given the findings that neither pH_i_ regulation nor ezrin interaction are the mechanism of action of NHE1-induced apoptosis resistance, we explored a third, non-canonical function of NHE1 as a plasma membrane scaffold for signaling pathways^32^. The cytoplasmic domain of NHE1 harbors binding sites for several signaling proteins^66^ including that of p38, and can modulate MAPK activity in response to diverse stimuli in other cell types^36^. Apoptotic stimulation with H_2_O_2_ increased p-38 phosphorylation in PASMCs from Nor rats as expected^46^ but not in PASMCs from SuHx animals. That NHE inhibition with EIPA and NHE1 silencing increased phosphorylation of p38 in PASMCs from SuHx rats supports the role of p38 phosphorylation in the NHE1-dependent regulation of apoptosis in SuHx PASMCs. Finally, we also confirmed the functional role of p38 phosphorylation in PASMC apoptosis as inhibition of p-p38 activity prevented the H_2_O_2_-induced increase in apoptosis in EIPA-treated SuHx PASMCs.

We speculate increased NHE1 expression in PASMCs from SuHx rats inhibits p38 phosphorylation by acting as a membrane-associated scaffold and sequestering signaling proteins through interactions mediated by its C-terminal cytoplasmic domain, although additional experimental evidence is needed to validate this hypothesis. These loss-of-function experiments were complemented by gain-function assays showing NHE1 wildtype and both mutants also prevented H_2_O_2_-induced p38 phosphorylation. Altogether, our results indicate repression of p38 phosphorylation is the mechanism by which NHE1 promotes resistance to apoptosis.

This study has several limitations. Although data from both male and female rats were included in all experiments, the limited number of female samples precluded formal sex-based analyses. Nonetheless, no apparent sex-related differences or trends were observed. We also noted discrepancies between Hoechst staining and caspase 3 activity results for some experiments. Possibilities for this discrepancy could be the late time point at which measurements were made, as caspase 3 activation is an early feature of apoptosis and/or can vary based on cell type and stimulus^67^. Future examination of caspase localization within the nucleus versus cytosol in PASMCs could also offer a more informative alternative to assessing total caspase activity, given that caspase translocation into the nucleus was shown to be required for apoptosis^68^. Lastly, the role of p-p38 in this study was examined via pharmacological inhibition with SB203580 and correlations with protein expression only. For future directions, replication of our gain-of-function assays with a mutation of the p38 binding motif in the cytosolic tail region of NHE1 to prevent the interaction between NHE1 and p38 as previously described^35^ could further define the role of p-p38 in NHE1-induced PASMC apoptosis resistance.

In summary, our results demonstrate NHE1 is required for the apoptosis-resistant phenotype of PASMCs from SuHx rats, and that its overexpression is sufficient to induce apoptosis resistance in control PASMCs (Fig 8). Furthermore, our findings indicate that modulation of p38 phosphorylation contributes to NHE1-medited apoptosis resistance independent of the roles of NHE1 in pHᵢ regulation or ezrin interaction. Collectively, these findings support the possibility that targeting NHE1-related pathways may help restore PASMC apoptosis and potentially reverse PH disease progression.

**Figure 8.**
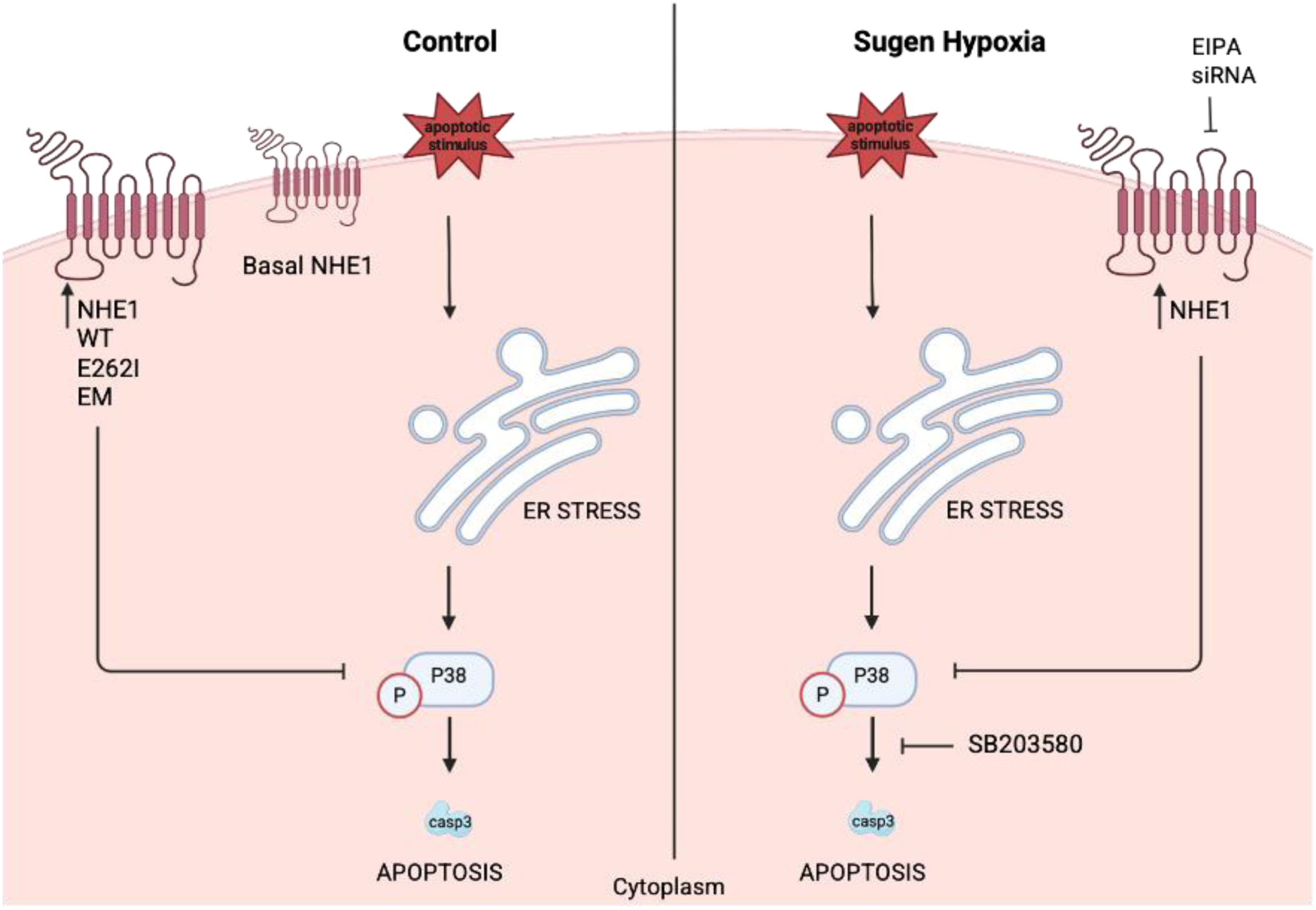
Schematic of proposed pathway. In control pulmonary arterial smooth muscle cells (PASMCs), apoptotic stimulation induces endoplasmic reticulum (ER) stress, activates p38 phosphorylation, and promotes apoptosis to maintain normal cell turnover. Overexpression of NHE1, including wild-type and mutant forms (E262l and EM), suppressed p38 phosphorylation and attenuated apoptosis. In PASMCs from Sugen/Hypoxia (SuHx) rats, elevated NHE1 surface expression and enhanced exchanger activity prevented p38 phosphorylation in response to apoptotic stimuli, thereby conferring apoptosis resistance. Pharmacologic inhibition of NHE1 with ethyl­isopropyl amiloride (EIPA) or gene silencing with siRNA restored p38 phosphorylation following apoptotic challenge and re-established apoptosis susceptibility. Conversely, inhibition of p38 activity using SB203580 blocked the pro-apoptotic effect of EIPA, confirming a key role for p38 signaling in NHEl-mediated apoptosis resistance in SuHx PASMCs.

## Bibliography

1. Kovacs G, Bartolome S, Denton CP, et al. Definition, classification and diagnosis of pulmonary hypertension. Eur Respir J. 2024;64(4):2401324. doi:10.1183/13993003.01324-2024

2. Humbert M, Kovacs G, Hoeper MM, et al. 2022 ESC/ERS Guidelines for the diagnosis and treatment of pulmonary hypertension: Developed by the task force for the diagnosis and treatment of pulmonary hypertension of the European Society of Cardiology (ESC) and the European Respiratory Society (ERS). Eur Heart J. 2022;43(38):3618-3731. doi:10.1093/eurheartj/ehac237

3. Shimoda LA, Laurie SS. Vascular remodeling in pulmonary hypertension. J Mol Med (Berl). 2013;91(3):297–309. doi:10.1007/s00109-013-0998-0

4. Humbert M, Guignabert C, Bonnet S, et al. Pathology and pathobiology of pulmonary hypertension: state of the art and research perspectives. Eur Respir J. 2019;53(1). doi:10.1183/13993003.01887-2018

5. Falcetti E, Hall SM, Phillips PG, et al. Smooth muscle proliferation and role of the prostacyclin (IP) receptor in idiopathic pulmonary arterial hypertension. Am J Respir Crit Care Med. 2010;182(9):1161–1170. doi:10.1164/rccm.201001-0011OC

6. Tajsic T, Morrell NW. Smooth muscle cell hypertrophy, proliferation, migration and apoptosis in pulmonary hypertension. Compr Physiol. 2011;1(1):295–317. doi:10.1002/cphy.c100026

7. Huetsch JC, Jiang H, Larrain C, Shimoda LA. The Na+/H+ exchanger contributes to increased smooth muscle proliferation and migration in a rat model of pulmonary arterial hypertension. Physiol Rep. 2016;4(5). doi:10.14814/phy2.12729

8. Paulin R, Meloche J, Courboulin A, et al. Targeting cell motility in pulmonary arterial hypertension. Eur Respir J. 2014;43(2):531–544. doi:10.1183/09031936.00181312

9. Leggett K, Maylor J, Undem C, et al. Hypoxia-induced migration in pulmonary arterial smooth muscle cells requires calcium-dependent upregulation of aquaporin 1. Am J Physiol Lung Cell Mol Physiol. 2012;303(4):L343–53. doi:10.1152/ajplung.00130.2012

10. Yun X, Niedermeyer S, Andrade MR, et al. Aquaporin 1 confers apoptosis resistance in pulmonary arterial smooth muscle cells from the SU5416 hypoxia rat model. Physiol Rep. 2024;12(16):e16156. 10.14814/phy2.16156

11. Benza RL, Miller DP, Barst RJ, Badesch DB, Frost AE, McGoon MD. An evaluation of long-term survival from time of diagnosis in pulmonary arterial hypertension from the REVEAL Registry. Chest. 2012;142(2):448–456. doi:10.1378/chest.11-1460

12. Bisserier M, Pradhan N, Hadri L. Current and emerging therapeutic approaches to pulmonary hypertension. Rev Cardiovasc Med. 2020;21(2):163–179. doi:10.31083/j.rcm.2020.02.597

13. Nicolls MR, Mizuno S, Taraseviciene-Stewart L, et al. New Models of Pulmonary Hypertension Based on VEGF Receptor Blockade-Induced Endothelial Cell Apoptosis. Pulm Circ. 2012;2(4):434–442. 10.4103/2045-8932.105031

14. Dromparis P, Sutendra G, Michelakis ED. The role of mitochondria in pulmonary vascular remodeling. J Mol Med (Berl). 2010;88(10):1003–1010. doi:10.1007/s00109-010-0670-x

15. Lampron M-C, Vitry G, Nadeau V, et al. PIM1 (Moloney Murine Leukemia Provirus Integration Site) Inhibition Decreases the Nonhomologous End-Joining DNA Damage Repair Signaling Pathway in Pulmonary Hypertension. Arterioscler Thromb Vasc Biol. 2020;40(3):783–801. doi:10.1161/ATVBAHA.119.313763

16. Meloche J, Le Guen M, Potus F, et al. miR-223 reverses experimental pulmonary arterial hypertension. Am J Physiol Cell Physiol. 2015;309(6):C363–72. doi:10.1152/ajpcell.00149.2015

17. Fuchs Y, Steller H. Programmed cell death in animal development and disease. Cell. 2011;147(4):742–758. doi:10.1016/j.cell.2011.10.033

18. Pan T, Zhang L, Miao K, Wang Y. A crucial role of endoplasmic reticulum stress in cellular responses during pulmonary arterial hypertension. Am J Transl Res. 2020;12(5):1481–1490.

19. Wu Y, Adi D, Long M, et al. 4-Phenylbutyric Acid Induces Protection against Pulmonary Arterial Hypertension in Rats. PLoS One. 2016;11(6):e0157538. doi:10.1371/journal.pone.0157538

20. Dromparis P, Paulin R, Stenson TH, Haromy A, Sutendra G, Michelakis ED. Attenuating Endoplasmic Reticulum Stress as a Novel Therapeutic Strategy in Pulmonary Hypertension. Circulation. 2013;127(1):115–125. doi:10.1161/CIRCULATIONAHA.112.133413

21. Quinn DA, Honeyman TW, Joseph PM, Thompson BT, Hales CA, Scheid CR. Contribution of Na+/H+ exchange to pH regulation in pulmonary artery smooth muscle cells. Am J Respir Cell Mol Biol. 1991;5(6):586–591. doi:10.1165/ajrcmb/5.6.586

22. Shimoda LA, Fallon M, Pisarcik S, Wang J, Semenza GL. HIF-1 regulates hypoxic induction of NHE1 expression and alkalinization of intracellular pH in pulmonary arterial myocytes. Am J Physiol Lung Cell Mol Physiol. 2006;291(5):L941–9. doi:10.1152/ajplung.00528.2005

23. Rios EJ, Fallon M, Wang J, Shimoda LA. Chronic hypoxia elevates intracellular pH and activates Na+/H+ exchange in pulmonary arterial smooth muscle cells. Am J Physiol Lung Cell Mol Physiol. 2005;289(5):L867–74. doi:10.1152/ajplung.00455.2004

24. Walker J, Undem C, Yun X, Lade J, Jiang H, Shimoda LA. Role of Rho kinase and Na+/H+ exchange in hypoxia-induced pulmonary arterial smooth muscle cell proliferation and migration. Physiol Rep. 2016;4(6). doi:10.14814/phy2.12702

25. Lade JM, Andrade MR, Undem C, et al. Hypoxia enhances interactions between Na(+)/H(+) exchanger isoform 1 and actin filaments via ezrin in pulmonary vascular smooth muscle. Front Physiol. 2023;14:1108304. doi:10.3389/fphys.2023.1108304

26. Quinn DA, Du HK, Thompson BT, Hales CA. Amiloride analogs inhibit chronic hypoxic pulmonary hypertension. Am J Respir Crit Care Med. 1998;157(4 Pt 1):1263–1268. doi:10.1164/ajrccm.157.4.9704106

27. Yu L, Quinn DA, Garg HG, Hales CA. Deficiency of the NHE1 gene prevents hypoxia-induced pulmonary hypertension and vascular remodeling. Am J Respir Crit Care Med. 2008;177(11):1276–1284. doi:10.1164/rccm.200710-1522OC

28. Yu L, Hales CA. Silencing of sodium-hydrogen exchanger 1 attenuates the proliferation, hypertrophy, and migration of pulmonary artery smooth muscle cells via E2F1. Am J Respir Cell Mol Biol. 2011;45(5):923–930. doi:10.1165/rcmb.2011-0032OC

29. Denker SP, Barber DL. Cell migration requires both ion translocation and cytoskeletal anchoring by the Na-H exchanger NHE1. J Cell Biol. 2002;159(6):1087–1096. doi:10.1083/jcb.200208050

30. Schelling JR, Abu Jawdeh BG. Regulation of cell survival by Na+/H+ exchanger-1. Am J Physiol Renal Physiol. 2008;295(3):F625–32. doi:10.1152/ajprenal.90212.2008

31. Wu KL, Khan S, Lakhe-Reddy S, et al. The NHE1 Na+/H+ exchanger recruits ezrin/radixin/moesin proteins to regulate Akt-dependent cell survival. J Biol Chem. 2004;279(25):26280–26286. doi:10.1074/jbc.M400814200

32. Baumgartner M, Patel H, Barber DL. Na(+)/H(+) exchanger NHE1 as plasma membrane scaffold in the assembly of signaling complexes. Am J Physiol Cell Physiol. 2004;287(4):C844–50. doi:10.1152/ajpcell.00094.2004

33. Wernig F, Mayr M, Xu Q. Mechanical Stretch-Induced Apoptosis in Smooth Muscle Cells Is Mediated by β1-Integrin Signaling Pathways. Hypertension. 2003;41(4):903–911. doi:10.1161/01.HYP.0000062882.42265.88

34. Gomez C, Martinez L, Mesa A, et al. Oxidative stress induces early-onset apoptosis of vascular smooth muscle cells and neointima formation in response to injury. Biosci Rep. 2015;35(4). doi:10.1042/BSR20140122

35. Grenier AL, Abu-ihweij K, Zhang G, et al. Apoptosis-induced alkalinization by the Na+/H+ exchanger isoform 1 is mediated through phosphorylation of amino acids Ser726 and Ser729. Am J Physiol Physiol. 2008;295(4):C883–C896. doi:10.1152/ajpcell.00574.2007

36. Pedersen SF, Darborg BV, Rentsch ML, Rasmussen M. Regulation of mitogen-activated protein kinase pathways by the plasma membrane Na+/H+ exchanger, NHE1. Arch Biochem Biophys. 2007;462(2):195–201. 10.1016/j.abb.2006.12.001

37. Yan S, Philip NM, Murray ST, et al. Physiological shear stress suppresses apoptosis in human pulmonary microvascular endothelial cells. Physiol Rep. 2025;13(6):e70269. 10.14814/phy2.70269

38. Sun X, Xue C, Jin Y, Bian C, Zhou N, Sun S. Glucose transporter GLUT1 expression is important for oriental river prawn (1. *Macrobrachium nipponense*) hemocyte adaptation to hypoxic conditions. J Biol Chem. 2023;299(1). doi:10.1016/j.jbc.2022.102748

39. Chen L, Mayer JA, Krisko TI, et al. Inhibition of the p38 kinase suppresses the proliferation of human ER-negative breast cancer cells. Cancer Res. 2009;69(23):8853–8861. doi:10.1158/0008-5472.CAN-09-1636

40. Banerjee M, Cui X, Li Z, et al. Na/K-ATPase Y260 Phosphorylation–mediated Src Regulation in Control of Aerobic Glycolysis and Tumor Growth. Sci Rep. 2018;8(1):12322. doi:10.1038/s41598-018-29995-2

41. Denker SP, Huang DC, Orlowski J, Furthmayr H, Barber DL. Direct binding of the Na--H exchanger NHE1 to ERM proteins regulates the cortical cytoskeleton and cell shape independently of H(+) translocation. Mol Cell. 2000;6(6):1425–1436. doi:10.1016/s1097-2765(00)00139-8

42. Fafournoux P, Noël J, Pouysségur J. Evidence that Na+/H+ exchanger isoforms NHE1 and NHE3 exist as stable dimers in membranes with a high degree of specificity for homodimers. J Biol Chem. 1994;269(4):2589–2596.

43. Quinn DA, Dahlberg CG, Bonventre JP, et al. The role of Na+/H+ exchange and growth factors in pulmonary artery smooth muscle cell proliferation. Am J Respir Cell Mol Biol. 1996;14(2):139–145. doi:10.1165/ajrcmb.14.2.8630263

44. Huetsch J, Shimoda LA. Na(+)/H(+) exchange and hypoxic pulmonary hypertension. Pulm Circ. 2015;5(2):228–243. doi:10.1086/680213

45. Son Y, Cheong Y-K, Kim N-H, Chung H-T, Kang DG, Pae H-O. Mitogen-Activated Protein Kinases and Reactive Oxygen Species: How Can ROS Activate MAPK Pathways? J Signal Transduct. 2011;2011(1):792639. 10.1155/2011/792639

46. McCubrey JA, LaHair MM, Franklin RA. Reactive Oxygen Species-Induced Activation of the MAP Kinase Signaling Pathways. Antioxid Redox Signal. 2006;8(9-10):1775–1789. doi:10.1089/ars.2006.8.1775

47. Riemann A, Schneider B, Ihling A, et al. Acidic environment leads to ROS-induced MAPK signaling in cancer cells. PLoS One. 2011;6(7):e22445. doi:10.1371/journal.pone.0022445

48. Simonin A, Fuster D. Nedd4-1 and &#x3b2;-Arrestin-1 Are Key Regulators of Na^+^/H^+^ Exchanger 1 Ubiquitylation, Endocytosis, and Function *. J Biol Chem. 2010;285(49):38293–38303. doi:10.1074/jbc.M110.115089

49. Ma Z, Yu Y-R, Badea CT, et al. Vascular Endothelial Growth Factor Receptor 3 Regulates Endothelial Function Through β-Arrestin 1. Circulation. 2019;139(13):1629–1642. doi:10.1161/CIRCULATIONAHA.118.034961

50. Santos CXC, Tanaka LY, Wosniak J, Laurindo FRM. Mechanisms and Implications of Reactive Oxygen Species Generation During the Unfolded Protein Response: Roles of Endoplasmic Reticulum Oxidoreductases, Mitochondrial Electron Transport, and NADPH Oxidase. Antioxid Redox Signal. 2009;11(10):2409–2427. doi:10.1089/ars.2009.2625

51. Hu Y, Yang W, Xie L, Liu T, Liu H, Liu B. Endoplasmic reticulum stress and pulmonary hypertension. Pulm Circ. 2020;10(1):2045894019900121. 10.1177/2045894019900121

52. Sehgal PB, Mukhopadhyay S. Dysfunctional intracellular trafficking in the pathobiology of pulmonary arterial hypertension. Am J Respir Cell Mol Biol. 2007;37(1):31–37. doi:10.1165/rcmb.2007-0066TR

53. Zhuan B, Wang X, Wang M-D, et al. Hypoxia induces pulmonary artery smooth muscle dysfunction through mitochondrial fragmentation-mediated endoplasmic reticulum stress. Aging (Albany NY). 2020;12(23):23684–23697. doi:10.18632/aging.103892

54. Stock C, Pedersen SF. Roles of pH and the Na+/H+ exchanger NHE1 in cancer: From cell biology and animal models to an emerging translational perspective? Semin Cancer Biol. 2017;43:5–16. 10.1016/j.semcancer.2016.12.001

55. Masereel B, Pochet L, Laeckmann D. An overview of inhibitors of Na(+)/H(+) exchanger. Eur J Med Chem. 2003;38(6):547–554. doi:10.1016/s0223-5234(03)00100-4

56. Karlsson M, Zhang C, Méar L, et al. A single–cell type transcriptomics map of human tissues. Sci Adv. 2025;7(31):eabh2169. doi:10.1126/sciadv.abh2169

57. Miyazaki E, Sakaguchi M, Wakabayashi S, Shigekawa M, Mihara K. NHE6 Protein Possesses a Signal Peptide Destined for Endoplasmic Reticulum Membrane and Localizes in Secretory Organelles of the Cell *. J Biol Chem. 2001;276(52):49221–49227. doi:10.1074/jbc.M106267200

58. Orlowski J, Grinstein S. Na+/H+ exchangers. Compr Physiol. 2011;1(4):2083–2100. doi:10.1002/cphy.c110020

59. Wei P-C, Lo W-T, Su M-I, Shew J-Y, Lee W-H. Non-targeting siRNA induces NPGPx expression to cooperate with exoribonuclease XRN2 for releasing the stress. Nucleic Acids Res. 2012;40(1):323–332. doi:10.1093/nar/gkr714

60. Betlej G, Błoniarz D, Lewińska A, Wnuk M. Non-targeting siRNA-mediated responses are associated with apoptosis in chemotherapy-induced senescent skin cancer cells. Chem Biol Interact. 2023;369:110254. 10.1016/j.cbi.2022.110254

61. Segal MS, Beem E. Effect of pH, ionic charge, and osmolality on cytochrome c-mediated caspase-3 activity. Am J Physiol Cell Physiol. 2001;281(4):C1196–204. doi:10.1152/ajpcell.2001.281.4.C1196

62. Lagadic-Gossmann D, Huc L, Lecureur V. Alterations of intracellular pH homeostasis in apoptosis: origins and roles. Cell Death Differ. 2004;11(9):953–961. doi:10.1038/sj.cdd.4401466

63. Parrish AB, Freel CD, Kornbluth S. Cellular mechanisms controlling caspase activation and function. Cold Spring Harb Perspect Biol. 2013;5(6). doi:10.1101/cshperspect.a008672

64. Dong Y, Gao Y, Ilie A, et al. Structure and mechanism of the human NHE1-CHP1 complex. Nat Commun. 2021;12(1):3474. doi:10.1038/s41467-021-23496-z

65. Dutta D, Fliegel L. Molecular modeling and inhibitor docking analysis of the Na+/H+ exchanger isoform one. Biochem Cell Biol. 2018;97(3):333–343. doi:10.1139/bcb-2018-0158

66. Slepkov ER, Rainey JK, Sykes BD, Fliegel L. Structural and functional analysis of the Na+/H+ exchanger. Biochem J. 2007;401(3):623–633. doi:10.1042/BJ20061062

67. Wolbers F, Buijtenhuijs P, Haanen C, Vermes I. Apoptotic cell death kinetics in vitro depend on the cell types and the inducers used. Apoptosis. 2004;9(3):385–392. doi:10.1023/B:APPT.0000025816.16399.7a

68. Del Rosario O, Suresh K, Kallem M, et al. MK2 nonenzymatically promotes nuclear translocation of caspase-3 and resultant apoptosis. Am J Physiol Cell Mol Physiol. 2023;324(5):L700–L711. doi:10.1152/ajplung.00340.2022

