## Supplemental figures for "Na^+^/H^+^ Exchanger Isoform 1 Regulates Apoptosis Susceptibility in Pulmonary Arterial Smooth Muscle from the Sugen/Hypoxia model of Pulmonary Hypertension"

**Supplemental Data**

Andrade et al.

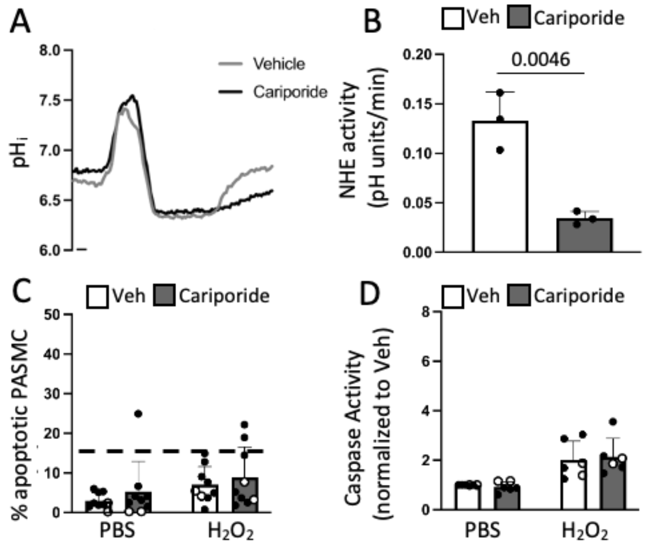


**Fig S1. Effects of $Na^+/H^+$ exchanger 1 (NHE1) inhibition with cariporide on apoptosis in Sugen/hypoxia (SuHx) pulmonary arterial smooth muscle cell (PASMC). A)** Representative traces and **B)** bar and scatter plots (mean +/- SD) show $Na^+/H^+$ exchanger (NHE) activity in PASMCs following treatment of cariporide (10 µM, 24 hr) or vehicle (DMSO) during the ammonium pulse protocol. Significance was assessed by unpaired two-tailed *t*-test. Bar and scatter plots show mean +/- SD for apoptosis in rat PASMCs from SuHx animals in response to stimulation with $H_20_2$ (250 µM, 24 hr) or vehicle (PBS) following cariporide incubation measured by **C)** Hoechst staining and **D)** caspase activity. Values for Hoechst staining are presented as percent of total cells while values for caspase activity are normalized to PBS and vehicle-treated SuHx cells. Significance assessed by two-way ANOVA with Holm-Sidak post hoc test. For **C)**, interaction p = 0.9009. For **D)**, interaction p = 0.8475.

Table 1: physiology of animal models

| | RV (g) | LV+S (g) | RVH | BW (g) |
|---|---|---|---|---|
| **Nor** | 0.204 + 0.027 | 0.774± 0.093 | 0.267 ±0.037 | 363.7 ± 54.5 |
| **SuHx** | 0.403 ± 0.066 | 0.780 ± 0.109 | 0.522 ± 0.078 | 295.1 ± 51.3 |
